# RNA fluorescence *in situ* hybridization (FISH) to visualize microbial colonization and infection in the *Caenorhabditis elegans* intestines

**DOI:** 10.1101/2022.02.26.482129

**Authors:** Dalaena E Rivera, Vladimir Lažetić, Emily R Troemel, Robert J Luallen

## Abstract

The intestines of wild *Caenorhabditis* nematodes are inhabited by a variety of microorganisms, including gut microbiome bacteria and pathogens, such as microsporidia and viruses. Because of the similarities between *Caenorhabditis elegans* and mammalian intestinal cells, as well as the power of the *C. elegans* system, this host has emerged as a model system to study host intestine-microbe interactions in vivo. While it is possible to observe some aspects of these interactions with bright-field microscopy, it is difficult to accurately classify microbes and characterize the extent of colonization or infection without more precise tools.

This protocol introduces RNA fluorescence in situ hybridization (FISH) as a tool used for the identification, visualization, and quantification of the microbes within the intestines of *C. elegans*. FISH probes that label the highly abundant small subunit ribosomal RNA can produce a bright signal for bacteria and microsporidian cells, and similar probes can be used to label viral RNA. FISH probes can be ordered from a commercial source as single-stranded DNA end-labeled with fluorophores. One limitation is that FISH may not provide robust signal against low copy targets, although signal can be boosted by using multiple probes (so-called ‘single-molecule FISH’). FISH staining involves collecting colonized or infected animals, washing to eliminate external contamination, followed by fixation in either paraformaldehyde or acetone. After fixation, FISH probes are incubated with samples to allow for the hybridization of probes to the desired target. To remove excess background, the animals are washed again, and then examined on microscope slides or using automated approaches.

Overall, this protocol enables detection, identification, and quantification of the microbes that inhabit the *C. elegans* intestine, including microbes for which there are no genetic tools available.

**SUMMARY:** Gut microbiome bacteria and intestinal intracellular pathogens, like the Orsay virus and microsporidia, are often found associated with wild *Caenorhabditis* nematodes. This protocol presents RNA FISH as a method for the detection, quantification, and identification of colonizing or infectious microbes within the context of intact *C. elegans* nematodes.

## INTRODUCTION

*Caenorhabditis elegans* has emerged as an excellent model system to study innate immunity and host-microbe interactions in the intestinal epithelial cells^1,2^. Due to having a transparent body and only 20 intestinal cells that are non-renewable, *C. elegans* represents a unique system for monitoring the processes of microbial intestinal colonization and infection in the context of an intact organism. Nematode intestinal cells share multiple morphological and functional similarities with mammalian intestinal epithelial cells making them a tractable in vivo model for dissection of conserved processes that govern microbiome colonization and pathogen infection^3,4,5,6^. *C. elegans* has additional advantages of a short life cycle and lifespan, and genetic tractability^7^.

Wild *C. elegans* feed on a variety of microbes that colonize and infect the intestine, and sampling of these nematodes has resulted in the discovery of viruses, fungi, oomycetes, and bacteria that naturally associate with this host^7,8,9,10^. The Orsay virus was found infecting the intestine and is currently the only known natural virus of *C. elegans*.^9^ Microsporidia are fungal-related obligate intracellular pathogens that are the most commonly found infection in wild caught *Caenorhabditis*, with several species having been discovered infecting *C. elegans* and related nematodes^8,11^. Many bacteria are commonly found inhabiting the intestinal lumen of wild-caught *C. elegans* and several species have been established as a natural model for the *C. elegans* microbiome (CeMbio)^6,12,13,14^. Discovering and characterizing microbes that naturally colonize and/or infect *C. elegans* is essential to understanding the genetic mechanisms that govern these host-microbe interactions, as well as visualizing novel microbial processes that only occur in the context of an intact host animal.

After sampling, wild nematodes are screened via differential interference contrast (DIC) microscopy to look for phenotypes that are indicative of infection or colonization. For example, changes in the stereotypical granulated appearance of the intestinal cells can be associated with the presence of an intracellular parasite infection^8^. Specifically, the loss of the gut granules and decreased cytosolic viscosity are signs of viral infection, whereas the reorganization of gut granules into ‘grooves’ may indicate infection with microsporidia in the genus *Nematocida*^8,9^. Because there is a wide variety of microbes present in a wild *C. elegans* samples, it can be difficult to distinguish among microbes through DIC microscopy. Information regarding the spatial distribution of microbes within the host may also be difficult to detect due to the small size of many microbes^15^. Additionally, culturing of any particular microbes of interest *in vitro* is not always possible, leading to difficulties in detection and/or quantification.

RNA fluorescence *in <u>s</u>itu* <u>h</u>ybridization (FISH) provides a method to fluorescently label microbes by utilizing fluorescent probes that bind to the RNA of the small ribosomal subunit in fixed cells. This protocol uses single-stranded DNA probes that are end-labeled with a fluorophore and specifically designed to be complementary to the target ribosomal sequence of the microbe of interest, although there are probes commercially available. The main advantage of targeting the small ribosomal subunit of microbes is the relatively large abundance of this RNA, leading to staining with a very high signal-to-noise ratio. Probes can also be designed to target genomic RNA to detect viruses, like the Orsay virus^9,16^. Some microbes can be identified and classified in a culture-independent manner using specific or universal FISH probes^8^. Additionally, RNA FISH can give insight on unique morphological colonization and infection characteristics, including filamentation or tissue localization patterns^17,18^. Different colored FISH probes can be used simultaneously, which allows for visual distinction between microbes in wild nematode samples, as well as observation of microbe-microbe dynamics inside a host^15,18^. Furthermore, infection and colonization can be easily quantified manually or through automated approaches^19^.

## PROTOCOL

1. Preparing nematodes with associated microbes
  1. Grow nematodes with the desired microbe of interest on standard Nematode Growth Media (NGM) plates seeded with the appropriate food source. Incubate the nematodes at 20°C until the desired life stage is reached.
  2. Add 2 mL M9 minimal salts media (42 mM Na2HPO4, 22 mM KH2PO4, 8.6 mM NaCl, 19 mM NH4Cl) + 0.1% Tween 20 to Nematode Growth Media (NGM) plates containing the *Caenorhabditis* strain infected or colonized with the desired microbe to be visualized. **NOTE:** 0.1% Triton-X can also be used in place of Tween 20 to make PBS-T.
  3. Pipette up the nematodes from the plates using a glass Pasteur pipette and bulb and transfer to 1.5 mL microfuge tubes. **NOTE:** Glass pipettes are preferred because nematodes can stick to plastic pipettes, but the addition of detergent (Tween 20 or Triton-X) can minimize this issue.
  4. Using a microcentrifuge, spin down the nematodes at 2000 x *g* for 60 s for L1’s or 500 x *g* for 60 s for L4 or adult animals. All subsequent centrifugation steps will be performed at the selected speed.
  5. Remove the supernatant from the microfuge tubes using a pipette. Avoid disturbing the nematode pellet by carefully removing the supernatant down to 100 µL above the pellet.
2. Washing nematodes to eliminate external contamination.
  1. Add 1 mL of 1x PBS (137 mM NaCl, 2.7 mM KCl, 10mM, Na2HPO4, 1.8 mM KH2PO4) + 0.1% Tween 20 (PBS-T) to microfuge tubes.
  2. Spin down the samples in microcentrifuge at the appropriate speed (see 1.3).
  3. Using a pipette, remove the supernatant. Leave approximately 100 µL of supernatant to avoid removing the pellet.
  4. Follow and repeat steps 2.1-2.3 two to three more times. **NOTE:** Three total washes are typically sufficient, however, additional washes can be performed to remove any excess bacteria.
3. Fixing nematodes **NOTE:** Nematodes may be fixed with either paraformaldehyde solution (PFA) or acetone. PFA allows for better visualization of morphology than acetone and can preserve signal from transgenic green fluorescent protein (GFP), which is destroyed by acetone. However, acetone fixation is necessary to permeabilize microsporidian spores to enable labeling this life stage. Furthermore, acetone can be more convenient than PFA because it is less toxic, and samples can be stored for several days in acetone in a -20 °C freezer without need for removing the fixative.
  1. To fix with PFA, in the fume hood, add 33 µL of 16% PFA to the microfuge tube containing 100 µL of supernatant above the nematode pellet obtained from step 2.3 for a final concentration of 4% PFA. **CAUTION:** PFA is a carcinogen. Contact with PFA may result in skin sensitization and irritation and eye damage. PFA releases toxic fumes that may lead to respiratory irritation or sensitization. When using PFA, work in a fume hood with proper personal protective equipment and refer to appropriate safety data sheets prior to use. **NOTE:** Alternatively, to fix with acetone, remove supernatant without disturbing the pellet and add 1 mL of acetone to the sample. **CAUTION:** Acetone is a highly flammable liquid and vapor. Although it can be purchased over the counter as nail polish remover, it is important to remember that acetone causes serious eye irritation and it may cause drowsiness or dizziness.
  2. For PFA fixation, incubate the nematode samples for 15-45 min at room temperature. **NOTE:** For acetone fixation, incubate the samples for 15 min at room temperature. **NOTE:** A shorter incubation period is better for maintaining GFP signal in transgenic strains due to degradation of GFP by PFA over time. Longer incubation times allow the fixative to better permeabilize into the samples. It is best to determine the incubation time empirically depending on the sample. **NOTE:** Following incubation, the samples can be stored until the protocol is ready to be continued. Samples fixed and incubated in PFA can be stored in 70% Ethanol at 4°C. Samples fixed and incubated in acetone can be stored in acetone at -20°C for up to two weeks.
4. Removing the fixative agent
  1. Spin down the nematode samples in a microcentrifuge at the appropriate speed (see 1.3).
  2. With a pipette, remove the supernatant without disturbing the pellet. **CAUTION:** The supernatant contains PFA, which is toxic. Discard the supernatant and at least the first two washes as toxic waste in a fume hood.
  3. Add 0.5 mL of PBS-T to the microfuge tubes.
  4. Follow and repeat steps 4.1-4.3 two to four times with PBS-T. **NOTE**: Performing more washes will help reduce background signal. Four washes in total are recommended.
  5. Following the last wash, spin down the samples and remove the supernatant, leaving the pellet undisturbed.
5. Preparing hybridization buffer (HB) and washing the nematodes
  1. Prepare 1 mL of HB (900 mM NaCl, 20 mM Tris pH 7.5, 0.01% SDS) per sample. Be sure to add SDS after water. **NOTE:** HB should be prepared fresh before each use to avoid precipitation. However, a general buffer (900 mM NaCl, 20 mM Tris pH 7.5) can be made in advance and stored at room temperature until HB is needed. Prior to use, prepare 1 mL per sample of the general buffer and add SDS to a final concentration of 0.01%.
  2. Add 800 µL of HB to the microfuge tubes containing the nematode pellet.
  3. Pellet the samples in the microcentrifuge. Remove the supernatant without disturbing the pellet.
6. Hybridizing FISH probe to desired target sequence
  1. Dilute FISH probe
    1. Mix 100 µL per sample of prepared HB with desired FISH probe to a final concentration of 5-10 ng/µL probe. **NOTE:** FISH probes are 15-23 mer oligos, antisense to the ribosomal subunit of the microbe of interest and labelled with a colored fluorophore attached to the 5’ or 3’ end (see **Table 1 for probes used here**).
  2. Add 100 µL of HB containing FISH probe to each sample. Mix by gently flicking or inverting the tubes. **NOTE:** Different colored FISH probes may be added simultaneously to visualize multiple fluorescent signals in the same sample (see Fig 1B).
  3. Incubate the samples overnight (6-24h) in a dry bath at 46-54°C or a thermal mixer at 46-54°C at 1200 rpm. **NOTE:** The temperature is dependent on the melting temperature (Tm) of the FISH probe and should be determined empirically.
7. Removing FISH probe and washing nematodes
  1. Prepare 3 mL of wash buffer (WB) (900 mM NaCl, 20 mM Tris pH 7.5, 5 mM EDTA, 0.01% SDS) per sample. **NOTE:** WB should be prepared fresh before each use to avoid precipitation. However, a general buffer (900 mM NaCl, 20 mM Tris pH 7.5) can be made in advance and stored at room temperature until wash buffer is needed. Prior to use, prepare 3 mL per sample of the general buffer and add EDTA to a final concentration of 5 mM and SDS to a final concentration of 0.01%.
  2. Centrifuge the samples at the appropriate speed (see 1.3). Remove the HB using a pipette, while being careful to leave the nematode pellet undisturbed.
  3. Add 1 mL of prepared WB to each sample.
  4. Centrifuge the samples at the appropriate speed (see 1.3). Remove the WB using a pipette, while being careful to leave the pellet undisturbed.
  5. Add 1 mL of prepared WB to each sample.
  6. Incubate the samples for 1 h at 48-56°C in a dry bath (or thermal mixer at 48-56°C at 1200 rpm). **NOTE:** The wash temperature is often 2°C higher than the hybridization temperature. **NOTE:** To further reduce background, incubation time in WB can be shortened to 30 min, after which, steps 7.4-7.5 should be repeated. This is to be followed by one (for bacteria) or two (for microsporidia sporoplasms) 30 min incubation periods at 46°C. **NOTE:** If incubating in a dry bath, gently invert the tubes approximately every 15-20 minutes.
  7. Centrifuge the samples at the appropriate speed (see 1.3). Using a pipette, remove the wash buffer, while being careful to leave the pellet undisturbed.
  8. Add 100-500 µL of PBS-T to each of the samples. **NOTE:** At this point, the samples can be stored in PBS-T at 4°C for up to a week until the protocol is ready to be continued.
8. Mounting the nematodes
  1. Pellet the samples at the appropriate speed (see 1.3). Remove as much PBS-T as possible without disturbing the nematode pellet.
  2. Add 20 µL of antifade mounting medium with DAPI (see **Table of Materials**) to the samples.
  3. Load a 20 µL pipettor with a 200 µL pipette tip and use scissors to cut the tip of the pipette off to allow larger nematodes to be pipetted.
  4. With the cut pipette tip, transfer 5-10 µL of the pellet onto a microscope slide. Cover with a 22×22 cover slip. **NOTE:** To store the slides, seal the edges of the cover slip with nail polish and keep in a dark box at 4°C until ready for further use.

**Table 1.**
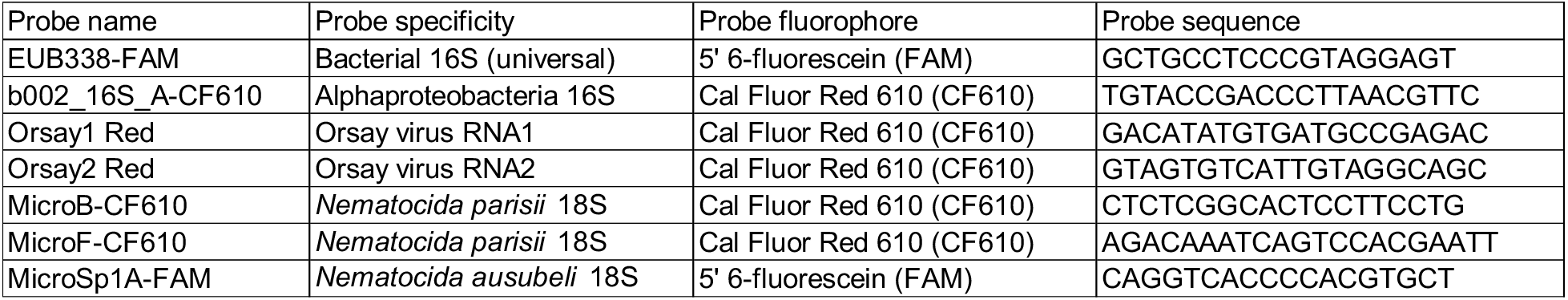
List of FISH probe sequences.

**Figure 1.**
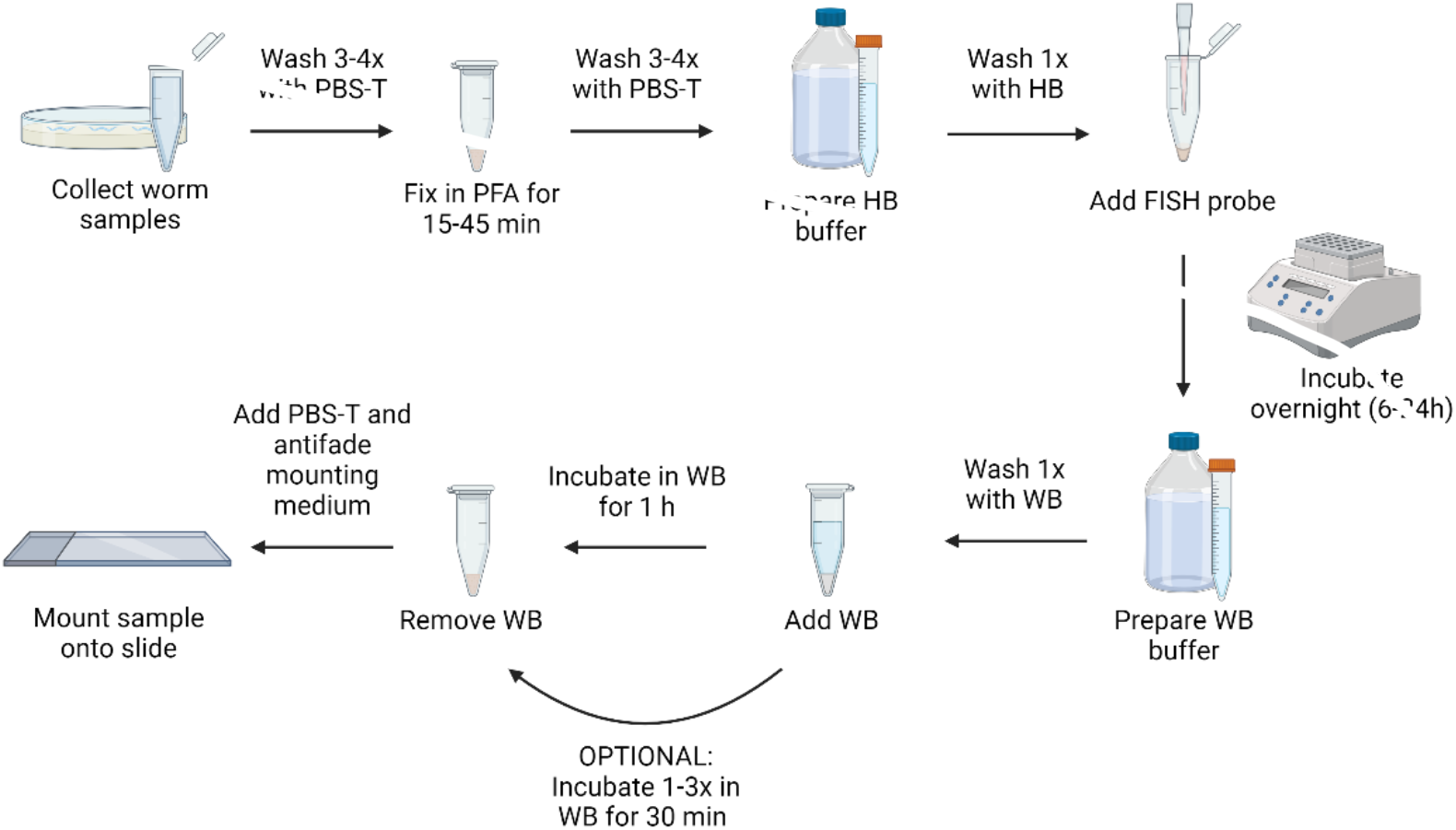
Visual representation of FISH protocol. Created with Biorender.com

## REPRESENTATIVE RESULTS

For looking at microbiome bacteria, specific and universal FISH probes to bacterial 16S were utilized on wild-isolated animals. Wild *Caenorhabditis tropicalis* strain (JU1848) was sampled from the Nouragues forest near a small river in the French Guiana from rotting palm tree fruits^20^. Under the DIC microscope, this nematode strain was found to be colonized with a bacterium that appears to directionally adhere to the intestinal epithelium (Fig. 2A). JU1848 was then selectively cleaned to eliminate other microbial contaminants and enrich for the desired adhering bacterium^21^. The bacterium was identified as a new species in the Enterobacteriaceae family. A FISH probe labeled with Cal Fluor Red 610 was designed specifically to the 16S rRNA sequence of this bacterium to allow fluorescent visualization of colonization within *C. tropicalis* (Fig. 2B). A universal 16S rRNA FISH probe capable of binding many species of bacteria (EUB338) was labeled with 6-carboxyfluorescin (FAM) and was also added to this sample. The green and red fluorescent signals overlap completely, suggesting the majority bacteria colonizing the intestines is the adhering *Enterobacteriaceae* bacterium. This bacterium causes distension in the lumen and colonizes nearly the entire anterior to posterior length of the worm.

**Figure 2.**
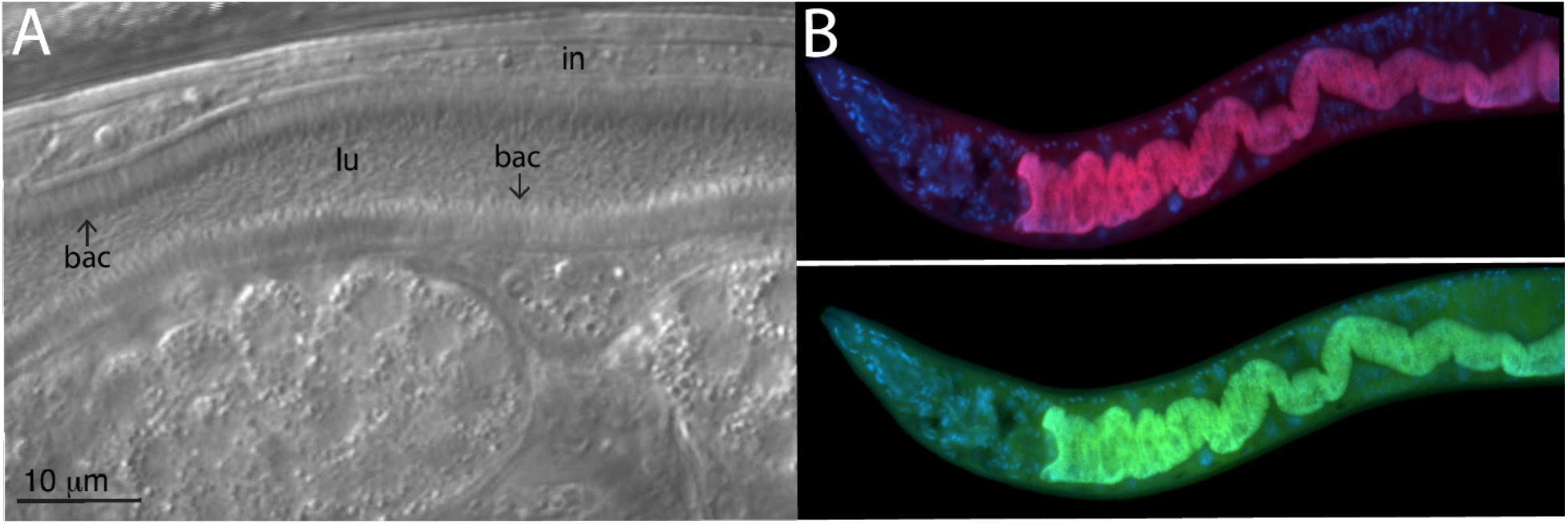
Wild *C. tropicalis* JU1848 strain colonized with adhering bacteria in the intestines. (A) Nomarski image depicting thousands of thin bacilli bacteria (ba) directionally binding to the intestine (in) of JU1848, creating a hair-like phenotype within the lumen (lu). This is adapted from Morgan, E., Longares, J. F., Félix, M. A., Luallen, R. J. Selective Cleaning of Wild *Caenorhabditis* Nematodes to Enrich for Intestinal Microbiome Bacteria. *J. Vis. Exp*. (174), e62937, doi:10.3791/62937 (2021)^21^. (B) FISH staining of JU1848 using a red labeled probe designed to the 16S rRNA sequence of the adhering bacterium (top) and a green labeled universal FISH probe to the 16S of bacteria (bottom). DAPI staining of host nuclei is shown in blue.

For looking at experimental infection with intracellular pathogens, Orsay virus and microsporidian-specific FISH probes were utilized on *C. elegans* with a wild-type background. The Orsay virus is a positive strand RNA virus from the *Nodaviridae* family, and the only natural viral pathogen found in *C. elegans*. The bipartite RNA genome of the Orsay virus consists of RNA1 and RNA2 segments, and FISH probes targeting both of these segments have been developed (Figures 3A and 3B)^9,16^. In the intestine, viral RNA is detected by RIG-I homolog DRH-1^22^, which is required for activation of the transcriptional defense program named the Intracellular Pathogen Response (IPR)^23,24,25^. The transcription of antiviral IPR genes is at least partially controlled by ZIP-1 transcription factor^19^. Here, the expression of ZIP-1::GFP is seen localized in the intestinal nuclei of cells that showing positive Orsay virus FISH staining in the cytoplasm (Figure 3A)^19^. Multiple animals stained with Orsay-specific FISH are shown to indicate the strength of this signal for easy quantification (Figure 3B).

**Figure 3.**
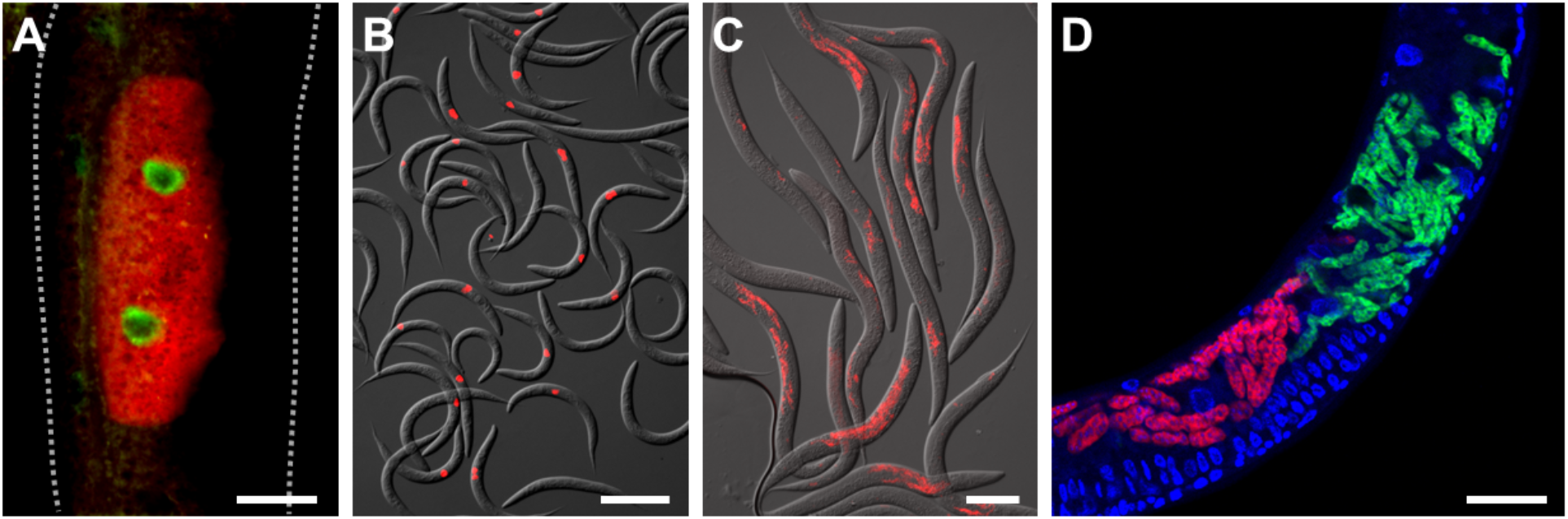
FISH staining of *C. elegans* infected with intracellular pathogens. (A, B) FISH staining of wild-type *C. elegans* infected with the Orsay virus. Orsay 1 Red and Orsay 2 Red probes were used for pathogen staining. (A) Composite image consists of merged red and green fluorescent channels. Nuclear ZIP-1::GFP expression is induced upon Orsay virus infection and is shown in green. Autofluorescence from the gut granules is shown in yellow. Dotted lines outline nematode body. Scale bar = 25 µm. (B) Composite image consists of merged red fluorescent and DIC channels. Scale bar = 200 µm. (C, D) FISH staining of wild-type *C. elegans* infected with microsporidia. (C) FISH staining of wild-type *C. elegans* infected with *N. parisii*. MicroB-CF610 probe was used for pathogen staining. Composite image consists of merged red fluorescent and DIC channels. Scale bar = 100 µm. (D) FISH staining of wild type *C. elegans* co-infected with *N. parisii* and *N. ausubeli* in the intestine. The two pathogens were co-stained using a pair of specific FISH probes that compete for binding to the same region of the 18S rRNA. *N. parisii* was stained using MicroF-CF610 (red) and *N. ausubeli* was stained using MicroSp1A-FAM (green). DAPI staining of host nuclei is seen in blue. Scale bar = 25 µm. (A-D) See Table 1 for probe sequences. Image shown in Figure 3A is adopted from: Lažetić, V., Wu, F., Cohen, L.B. et al. The transcription factor ZIP-1 promotes resistance to intracellular infection in *Caenorhabditis elegans. Nat Commun*. **13**, 17. (2022). https://doi.org/10.1038/s41467-021-27621-w^19^. The changes were made to the original published image. Creative Commons license: https://creativecommons.org/licenses/by/4.0/ Images shown in Figure 3B and 3C are adopted from: Reddy KC, Dror T, Underwood RS, Osman GA, Elder CR, Desjardins CA, et al. (2019). Antagonistic paralogs control a switch between growth and pathogen resistance in *C. elegans. PLoS Pathog*. **15** (1), e1007528. https://doi.org/10.1371/journal.ppat.1007528^24^. The changes were made to the original published image. Creative Commons license: https://creativecommons.org/licenses/by/4.0/

The microsporidian parasite named *Nematocida parisii*, meaning nematode killer from Paris, is an obligate intracellular pathogen of the intestine. Several FISH probes that label 18S rRNA of *N. parisii* have been used, including fluorescently tagged MicroB and MicroF probes. Multiple animals stained with MicroB FISH are shown to indicate the strength of this signal for easy quantification (Figure 3C). Additionally, *C. elegans* is infected by other closely related microsporidia. Co-infection of N2 with *N. parisii* and the related *N. ausubeli* can be distinguished using this FISH protocol by designing species-specific FISH probes that compete for the same binding site on the 18S rRNA (Figure 3D)^26^. Additionally, the use of DAPI to stain nuclei allow for better localization of the infection in the context of the whole animal, especially for the intestine which have large, easily identifiable nuclei.

## DISCUSSION

Wild *C. elegans* are naturally associated with a variety of microbes. Researchers can use RNA FISH to detect and identify these microbes as well as gain insight into their localization in the context of a whole animal. The abundance of numerous bacterial isolates from wild *C. elegans* may also be quantified via RNA FISH^27^. By using the protocol described here, it is also possible to observe these microorganisms inside their hosts and learn more about their interactions. Importantly, Orsay virus and microsporidia are obligate parasites and cannot be cultured independently of the host, and there are no genetically modified versions of pathogens that express a fluorescent protein, so FISH is the standard visualization tool. In addition to staining microorganisms in *C. elegans* intestine, this protocol can be used for other nematode strains like *C. tropicalis* or *Oscheius tipulae*^17,21^.

The main advantages of this protocol are that it offers a simple, quick, and robust method to stain for microbes associated with *C. elegans*. Images produced from FISH have a high signal-to-noise ratio, which is achieved by utilizing FISH probes that target the abundant small subunit ribosomal RNA within the sample. Furthermore, RNA FISH makes it possible to see infection or colonization within the context of the whole animal. This visualization is facilitated through co-staining host nuclei with DAPI and/or using fluorescent-marked strains of *C. elegans* to better highlight the localization of infection or colonization within the sample. For example, microsporidian-specific FISH was used to determine the tissue tropism of *Nematocida displodere* by using a panel of *C. elegans* strains with GFP expression in different tissues^18^. Additionally, this protocol is amenable to changes that allow for researchers to determine the ideal conditions suitable for their specific needs, e.g., adjusting the fixation period, increasing the hybridization temperature.

One critical step in this protocol is fixing the samples. The incubation period following the addition of the fixative is necessary to allow time for the agent to permeabilize the sample. Longer incubation times are not ideal for samples containing transgenic fluorescent proteins due to protein degradation by PFA over time. For samples containing GFP, it is imperative to determine the optimal fixation time to allow for permeabilization, while still maintaining GFP signal.

FISH can be used to stain for bacteria, viruses, or microsporidia in *C. elegans*. However, the best type of fixative agent used for FISH depends on the sample and downstream requirements. This protocol presents a PFA solution as the primary fixative agent to stain for bacteria and viruses. However, PFA is not sufficient for the visualization of microsporidian spores as it cannot penetrate the spore wall. For visualization of spores, acetone should be used instead. Although, PFA fixation is efficient for FISH labeling of other life stages of microsporidia, including sporoplasms, meronts, and sporonts. We see other major differences between using acetone and PFA fixation. Acetone is more convenient because samples can be quickly stored in the freezer after adding, without need for washing. However, acetone quickly kills any existing GFP in the host. PFA is the preferred fixative if it is important to preserve some physiological structures in the host, as acetone-fixed animals appear to be more degraded, making identification of some tissues more difficult. Because the samples are fixed, this FISH protocol does not allow for live imaging of host-microbe interactions *in vivo*. However, a pulse-chase infection time course followed by FISH staining samples at various timepoints can allow one to see some dynamics of microbial infection^17,18,28^.

Another critical step throughout the protocol is thoroughly washing the samples before and after hybridization. Before hybridization, when collecting the worms into the microfuge tubes, excess bacteria or other microbes from the NGM plates can be carried with the worm sample. Three washes with PBS-T are standard, however, more washes may be necessary to sufficiently eliminate external microorganisms, especially when using heavily contaminated, wild-isolated *C. elegans*. When viewing the mounted samples after FISH, there may be some residual FISH probe that produces large amounts of signal in the background of the sample. The wash temperature and the number of washes is important to remove the excess and non-specifically bound probe. To reduce the background fluorescence, it is possible to perform two or three washes with 1 mL of WB every 30 minutes, instead of one wash with 1 mL of WB for an hour. Different FISH probes may require different wash temperatures. Typically, the wash temperature is 2°C above the hybridization temperature, but this can be increased if there is too much background fluorescence (high noise).

This protocol utilizes fluorescent probes that target microbial rRNA, but FISH probes can be designed to other high-copy transcripts. Other FISH probes may have different melting temperatures, so incubation steps may need to be performed at a higher or lower temperature than described. This FISH protocol can identify the spatial distribution of microbial colonization or infection within the host, allowing for the characterization of host-microbe and microbe-microbe interactions. One limitation to this protocol is that some microorganisms cannot be phylogenetically distinguished from one another simultaneously via different colored fluorophores. This reduces the number of different microorganisms that can be detected via FISH at the same time which limits its use for complex microbiome studies in *C. elegans*. However, multicolor rRNA-targeted FISH utilizes probes labeled with non-canonical fluorophores can increase the number of distinct microbial group labels^15^. Another limitation is that this protocol cannot distinguish between closely related species, especially bacteria, that have ribosomal RNA sequences that are highly similar. However, the extreme sequence divergence between microsporidia species helps to facilitate their differentiation with this protocol (Fig. 3)^29,30^.

Overall, this FISH protocol describes a technique to detect microorganisms within *C. elegans*. This protocol allows researchers to use a transparent and genetically tractable model system to detect and quantify colonization and infection *in vivo*, as well as identify unique microbial behavior or morphology within the host.

## ACKNOWLEDGMENTS

Thank you to Dr. Marie-Anne Félix for providing us with wild nematode strains.

## REFERENCES

1. Pukkila-Worley, R., Ausubel, F. M. Immune defense mechanisms in the Caenorhabditis elegans intestinal epithelium. Current Opinion in Immunology. 24, 3–9 (2012)

2. Balla, K. M., Troemel, E. R. Caenorhabditis elegans as a model for intracellular pathogen infection. Cellular Microbiology. 15, 1313–1322 (2013)

3. Dimov, I., Maduro, M. F. The C. elegans intestine: organogenesis, digestion, and physiology. Cell and Tissue Research. 377, 383–396 (2019).

4. Bossinger, O., Fukushige, T., Claeys, M., Borgonie, G., McGhee, J. D. The apical disposition of the Caenorhabditis elegans intestinal terminal web is maintained by LET-413. Developmental Biology. 268, 448–456 (2004).

5. Szumowski, S. C., Botts, M. R., Popovich J. J., Smelkinson M. G., Troemel, E. R. The small GTPase RAB-11 directs polarized exocytosis of the intracellular pathogen N. parisii for fecal-oral transmission from C. elegans. PNAS. 111 (22), 8215–8220 (2014).

6. Samuel, B.S., Rowedder, H., Braendle, C., Félix, M.A., Ruvkun, G. Caenorhabditis elegans responses to bacteria from its natural habitats. PNAS. 113 (27), E3941–E3949 (2016).

7. Zhang, F. et al. Caenorhabditis elegans as a model for microbiome research. Frontiers in Microbiology. 8, 485 (2017)

8. Troemel, E. R., Félix, M.-A., Whiteman, N. K., Barrière, A., Ausubel, F. M. Microsporidia are natural intracellular parasites of the nematode Caenorhabditis elegans. PLoS Biology. 6, 2736–2752 (2008).

9. Felix, M.A. et al. Natural and experimental infection of Caenorhabditis nematodes by novel viruses related to nodaviruses. PLoS Biol. 9, e1000586 (2011).

10. Osman, G.A., et al. Natural Infection of C. elegans by an Oomycete Reveals a New Pathogen-Specific Immune Response. Curr Biol. 28 (4), 640–648 (2018).

11. Zhang, G. A Large Collection of Novel Nematode-infecting Microsporidia and their Diverse Interactions with Caenorhabditis elegans and other related nematodes. PLOS Path. 12 (12) e1006093. (2016)

12. Clark, L.C., Hodgkin, J. Commensals, probiotics and pathogens in the Caenorhabditis elegans model. Cell Microbiology. 16, 27–38 (2014).

13. Dirksen, P. et al. CeMbio - The Caenorhabditis elegans Microbiome Resource. G3 (Bethesda). 10 (9), 3025–3039 (2020).

14. Berg, M., Stenuit, B., Ho, J. et al. Assembly of the Caenorhabditis elegans gut microbiota from diverse soil microbial environments. ISME J 10, 1998–2009 (2016).

15. Michael, L., Markus, S., Petra, Pjevac., Holger, D. A Multicolor Fluorescence in situ Hybridization Approach Using an Extended Set of Fluorophores to Visualize Microorganisms. Frontiers in Microbiology. 10, 1383 (2019).

16. Franz, C.J. et al. Orsay, Santeuil and Le Blanc viruses primarily infect intestinal cells in Caenorhabditis nematodes. Virology. 448, 255–264. (2014)

17. Tran, T.D., Ali, M.A., Lee, D. et al. Bacterial filamentation as a mechanism for cell-to-cell spread within an animal host. Nat Commun 13, 693 (2022).

18. Luallen, R.J. et al. Discovery of a natural microsporidian pathogen with broad tissue tropism in Caenorhabditis elegans. PLOS Pathogens. 12 (6), e1005724 (2016).

19. Lažetić, V. et al. The transcription factor ZIP-1 promotes resistance to intracellular infection in Caenorhabditis elegans. Nat Commun. 13, 17 (2022).

20. Félix, M. -A., et al. Species richness, distribution and genetic diversity of Caenorhabditis nematodes in a remote tropical rainforest. BMC Evolutionary Biology. 13, 10 (2013).

21. Morgan, E., Longares, J.F., Félix, M.A., Luallen, R.J. Selective Cleaning of Wild Caenorhabditis Nematodes to Enrich for Intestinal Microbiome Bacteria. JoVE. (174), e62937 (2021).

22. Sowa, J. N. et al. The Caenorhabditis elegans RIGI Homolog DRH-1 Mediates the Intracellular Pathogen Response upon Viral Infection. Journal of Virology. 94, e01173–19 (2020).

23. Bakowski, M.A. et al. Ubiquitin-Mediated Response to Microsporidia and Virus Infection in C. elegans. PLOS Pathogens. 10 (6), e1004200. (2014).

24. Reddy, K.C. et al. Antagonistic paralogs control a switch between growth and pathogen resistance in C. elegans. PLoS Pathog. 15 (1), e1007528. (2019).

25. Reddy, K.C. et al. An Intracellular Pathogen Response Pathway Promotes Proteostasis in C. elegans. Curr Biol. 27 (22), 3544–3553. (2017).

26. Balla, K.M., Lažetić, V., Troemel, E.R. Natural variation in the roles of C. elegans autophagy components during microsporidia infection. PloS one, 14 (4), e0216011. (2019).

27. Dirksen, P., et al. The native microbiome of the nematode Caenorhabditis elegans: gateway to a new host-microbiome model. BMC Biol 14, 38 (2016).

28. Willis, A.R., et al. A parental transcriptional response to microsporidia infection induces inherited immunity in offspring. Science Advances. 7 (19). (2021).

29. Cuomo, C.A., et al. Microsporidian genome analysis reveals evolutionary strategies for obligate intracellular growth. Genome Res. 22 (12), 2478–88. (2012).

30. Reinke, A.W., Balla, K.M., Bennett, E.J., Troemel, E.R. Identification of microsporidia host-exposed proteins reveals a repertoire of rapidly evolving proteins. Nature Commun. 8, 14023. (2017).

